# An investigation on the feasibility of shipping protein crystals in plates for room temperature *in situ* data collection

**DOI:** 10.1101/2025.06.12.659325

**Authors:** Christopher S. Campomizzi, M. Elizabeth Snell, Halina Mikolajek, James Sandy, Juan Sanchez-Weatherby, Amy J. Thompson, Gabrielle R. Budziszewski, Silvia Russi, Richard Howells, Hannah Walters-Morgan, Ralf Flaig, Aina Cohen, Michael A. Hough, Sarah E.J. Bowman

**Affiliations:** Department of Biochemistry, Jacobs School of Medicine and Biomedical Sciences, University at Buffalo, 955 Main Street, Buffalo, NY, 14203, United States; University at Buffalo Hauptman Woodward Institute, University at Buffalo, Buffalo, NY, 14203, United States; Diamond Light Source Ltd, Harwell Science and Innovation Campus, Didcot, OX11 0DE, United Kingdom; Research Complex at Harwell, Harwell Science and Innovation Campus, Didcot, OX11 0FA, United Kingdom; Stanford Synchrotron Radiation Lightsource, SLAC National Accelerator Laboratory, Stanford University, Menlo Park, CA, 94025, United States; Crystal Positioning Systems, 1070 Allen Street, Jamestown, NY, 14701, United States

**Keywords:** Room-temperature crystallography, *in-situ* X-ray diffraction, protein crystal shipping

## Abstract

Room-temperature X-ray diffraction experiments can facilitate investigation of protein dynamics, efficient probing of fragment binding, and time-resolved crystallography experiments. The Versatile Macromolecular Crystallography *in situ* (VMXi) beamline at Diamond Light Source (DLS) in the United Kingdom specializes in the collection of room-temperature X-ray diffraction data *in situ* directly from crystallization plates without any manipulation of protein crystals, preserving the integrity of fragile samples. While many X-ray light sources are now equipped to grow crystals on site for room-temperature experiments, to date there has been no comprehensive analysis of the effect of shipping crystals within plates at ambient temperature for *in situ* room-temperature data collection. *In situ* methods place more stringent demands on shipping, as crystallization drops must arrive both intact and properly positioned within the plate to remain in the field of view of crystal imaging systems. The latter requirement imposes more stringent limits on shipping procedures as lateral movement of <1mm may leave a droplet out of the crystal imager field of view yet is independent of the requirement for the diffraction quality of crystals to be maintained. In contrast, the equivalent methodology for cryo-cooled crystals is well established. Here we investigate the impact of transatlantic shipping of crystals generated from commercially available proteins (lysozyme, thaumatin, and thermolysin) within the same plates in which they were grown. MiTeGen In Situ-1™ plates containing the protein crystals were set up at the University of Buffalo Hauptman Woodward Research Institute (UB-HWI, Buffalo, NY, United States) and shipped to DLS (Didcot, United Kingdom). Identical plates were prepared on site at DLS in the Crystallisation Facility at Harwell. We also performed this experiment with human lysine acetyltransferase 2B (KAT2B) to validate our proposed pipeline with a typical protein crystal sample. For shipping, we utilized the Stanford Synchrotron Radiation Lightsource (SSRL) Blue Thermal Shipping Box (Blue Box), which can maintain temperature for a few days, to ship crystallization plates at room temperature from UB-HWI to DLS. We hypothesized that long-distance shipping by standard commercial courier might compromise data quality through mechanical stress or temperature fluctuations. Instead, we found that room-temperature data collected at VMXi showed no significant differences for crystals set up at UB-HWI and shipped relative to crystals set up on site in the UK. High-resolution structures were successfully determined for all proteins in the study (both those shipped from UB-HWI and set up on-site at DLS), demonstrating that long-distance shipment of crystals at non-cryogenic temperatures is feasible without compromising diffraction quality. This study provides a proof-of-concept workflow for expanding access to room-temperature crystallography worldwide, enabling more researchers to leverage cutting-edge techniques without needing to grow crystals on site.

**Synopsis:** The effect of international shipment of protein crystals on *in situ* room-temperature X-ray diffraction quality.

## 1. Introduction

X-ray crystallography remains the predominant technique for the determination of three-dimensional biomolecular structures, accounting for over 80% of structures deposited into the Protein Data Bank (https://www.rcsb.org/stats/summary) (Berman *et al*., 2000). While methods such as cryo-electron microscopy (Cryo-EM) are becoming more prevalent for structure determination, it was demonstrated during the COVID-19 pandemic that both X-ray crystallography and Cryo-EM provide important structural information (Lynch *et al*., 2021). Additionally, diffraction-based methods are currently undergoing an explosion of technique development (Ebrahim *et al*., 2022, Moreno-Chicano *et al*., 2022, Okumura *et al*., 2022, Bjelcic *et al*., 2023, Mikolajek *et al*., 2023, Foos *et al*., 2024, Khusainov *et al*., 2024) enabled by advances in X-ray sources and beamline optics (McNeil & Thompson, 2010, Chapman *et al*., 2011, Eriksson *et al*., 2014, Sierra *et al*., 2019, Schneider *et al*., 2021, Warren *et al*., 2024, Hegde *et al*., 2025), as well as detector technology (Dinapoli *et al*., 2011, Hatsui & Graafsma, 2015, Mozzanica *et al*., 2018, Tolstikova *et al*., 2019, van Driel *et al*., 2020).

An overwhelming majority of structures from X-ray crystallography have been determined at cryogenic (cryo) temperature. Cryocrystallography has contributed extensively to structural biology, offering several practical advantages, including a substantial increase in the X-ray dose a sample can withstand before experiencing significant diffraction loss due to radiation damage (Young *et al*., 1990), as well as ease of transport, and compatibility with remote data collection (Soltis *et al*., 2008). Despite these advantages, there are limitations in using cryo-cooled crystals. First, cryo-cooling a crystal can ‘trap’ the macromolecule in a non-biologically relevant protein conformation. Second, protein crystals are often delicate, making them difficult to harvest and cryopreserve (Teng, 1990). Third, structures from diffraction data collected at cryogenic temperatures below the glass transition are snapshots of a particular state of a protein; dynamics and conformational fluctuations are not typically observable(Juers & Matthews, 2001, Keedy *et al*., 2014, Keedy, Fraser*, et al.*, 2015).

Room-temperature (RT) crystallography is re-emerging as a valuable approach. While it may not compete with techniques like cryocrystallography and Cryo-EM for solving novel structures, room-temperature crystallography offers insight into structural data that often aligns better with data collected from solution-based methods, which in many cases are more biologically relevant (Fraser *et al*., 2009, Keedy, Kenner*, et al.*, 2015, Fenwick *et al*., 2014). Room-temperature crystallography can therefore offer insight into protein dynamics and ligand interactions. Furthermore, there is no need for the addition of cryoprotectants, which may disrupt local structure and ligand binding sites. Notably, the field of time-resolved serial crystallography at X-ray Free Electron Laser sources and synchrotron beam lines is almost entirely dependent on room-temperature crystallographic experiments (Chapman *et al*., 2011, Gati *et al*., 2014).

Despite its potential, room-temperature crystallography presents several key challenges, particularly in crystal transportation and handling. While shipping of dewars containing cryo-cooled crystals has become routine with most synchrotron users using this approach together with remote access control of X-ray beamlines or fully automated unattended data collection, equivalent approaches for room-temperature crystals are far less well developed. One major challenge is keeping crystals hydrated during transportation and sample handling at the beamline. Originally, crystals were frequently transported by hand in polystyrene boxes and mounted in glass capillaries (Blundell & Johnson, 1976); the use of plastic capillaries has also been implemented with success (Kalinin *et al*., 2005). Controlled dehydration using hydration streams at beamlines has been used for improving resolution (Sanchez-Weatherby *et al*., 2009, Russi *et al*., 2011). Another major challenge is the decrease in diffraction lifetime, as radiation damage is more pronounced at non-cryogenic temperatures. To address this challenge, data from multiple crystals can be collected and combined into a single dataset (Vaughan *et al*., 2004, Gildea *et al*., 2022, Thompson *et al*., 2025). While shipping crystals is common for cryocrystallography, most samples for room-temperature crystallography, including those grown for *in situ* experiments, are grown at the X-ray source (Bingel-Erlenmeyer *et al*., 2011, le Maire *et al*., 2011, Axford *et al*., 2012). In fact, to our knowledge, only one published study exists detailing a means to ship room-temperature crystals, in this case for a membrane protein grown using *in meso* techniques (Huang *et al*., 2015).

The Versatile Macromolecular Xtallography *in situ* (VMXi) beamline at Diamond Light Source (DLS) in the United Kingdom specializes in the highly automated collection of room-temperature X-ray diffraction data directly from 96-well crystallization plates, with the recommended plate type being MiTeGen In Situ-1™ (Sanchez-Weatherby *et al*., 2019, Mikolajek *et al*., 2023). The *in situ* plate-based data collection method provides two major benefits for collecting room-temperature crystallographic data: it eliminates the need for physical manipulation of fragile crystals and reduces the risk of dehydration by maintaining a sealed environment. Prior to our experiments, there were three modes of access available to users at VMXi: 1) users could travel with their pre-dispensed plates to the beamline, 2) users could travel to the crystallization facility, which is part of the Research Complex at Harwell, to set up plates, or 3) users could ship protein samples to the crystallization facility to be setup at DLS. Currently, the majority of VMXi users are from within the United Kingdom; while there are international users, the need for on-site crystallization previously limited accessibility. While the shipping of crystals for cryocrystallography at 100 K in shipping dewars has become standardized and reliable, robust standardized methods to ship room-temperature crystals for *in situ* data collection have not yet been established.

Here, we present results showing the successful shipping of protein crystals, grown on MiTeGen In Situ-1™ plates, from the University at Buffalo Hauptman Woodward Research Institute (UB-HWI) in Buffalo, New York, USA to DLS in Didcot, UK. We used the Stanford Synchrotron Radiation Lightsource (SSRL) Blue Thermal Shipping Box (Blue Box), which can maintain temperature for several days, to ship crystallization plates at room-temperature from UB-HWI to DLS (Figure 1). The Blue Box was initially intended for use with SSRL *in situ* plates, and together these components are integral to a general user program at SSRL that supports fully remote-access data collection using crystals at controlled non-cryogenic temperatures and controlled-humidity conditions at SSRL beamlines 12-1 and 9-2. While developed to accommodate SSRL *in situ* plates, to facilitate remote collection at other facilities, the SSRL Blue Box was also designed to accommodate standard crystallization plates. Our goals were to investigate whether shipping room-temperature crystals long distances, in this case internationally, using a standard shipping option from a commercial shipping company, has any impact on data quality, and whether shipping crystals grown directly on MiTeGen In-Situ plates to VMXi is feasible. Identical crystallization plates were set up at UB-HWI and DLS with the well-characterized proteins lysozyme, thaumatin, and thermolysin, in order to compare the quality of diffraction and structures from crystals grown at each location. Our aim was to collect high-quality, room-temperature crystallographic data at VMXi for the crystals shipped from UB-HWI that could be compared to data from samples set up on-site at DLS. To verify that our pipeline is effective for typical protein crystals, rather than only for robust crystals of standard proteins, we also collected similar data on human lysine acetyltransferase 2B (KAT2B), an important DNA damage response protein (Sakaguchi *et al*., 1998, Liu *et al*., 1999) with and without return shipment between the UK and USA.

**Figure 1.**
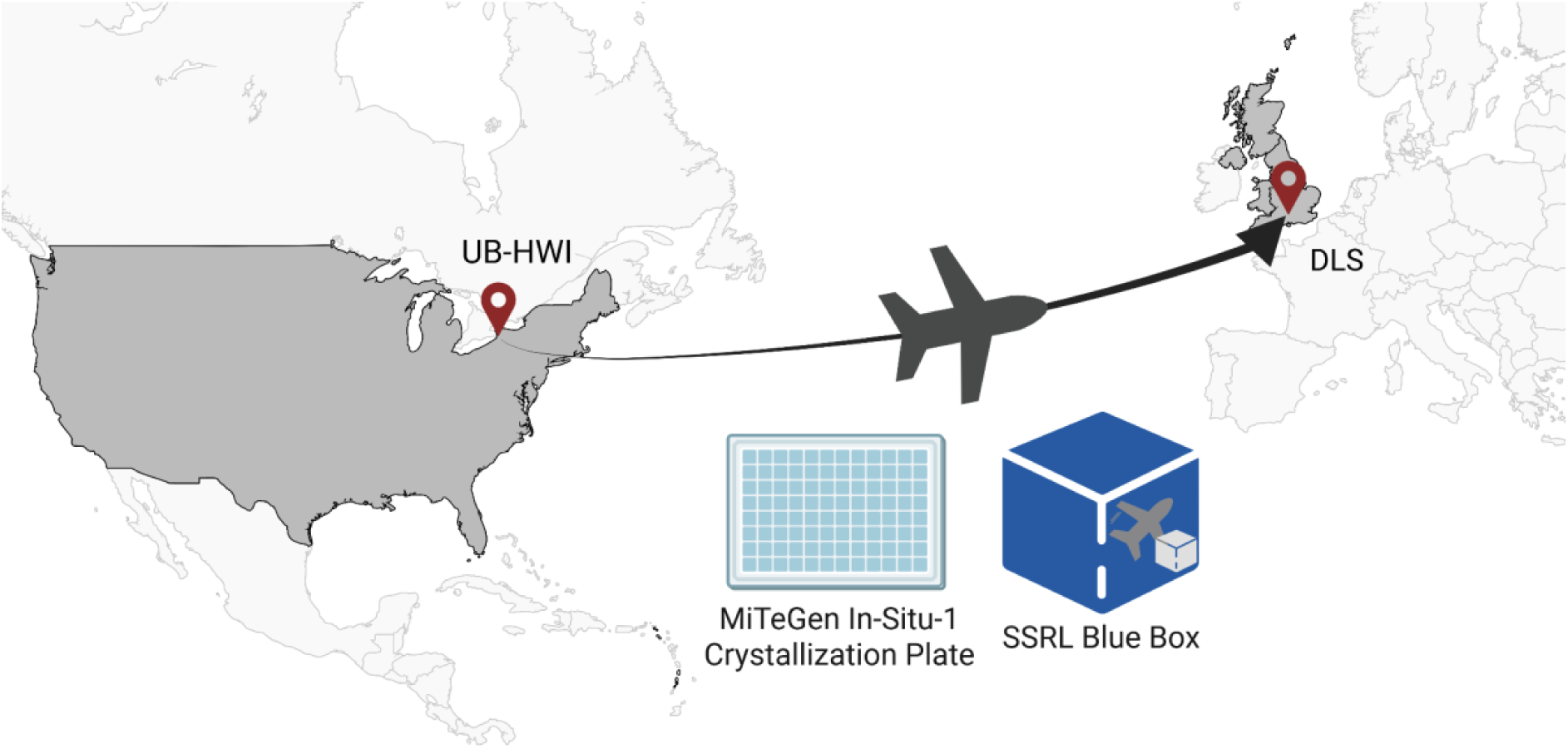
Schematic overview of the experimental workflow. Samples were grown on MiTeGen In Situ-1™ crystallization plates at UB-HWI, packed into the SSRL Blue Box, and transported by air and road approximately 3500 miles (5650 kilometers) via standard commercial shipping to DLS in the United Kingdom for room-temperature, *in situ* X-ray diffraction experiments. Note that for experiments with KAT2B, crystals in plates were shipped from DLS to HWI, imaged on site and then shipped back to DLS for data collection

We hypothesized that long-distance, room-temperature shipping of protein crystals might compromise data quality due to mechanical shock or thermal fluctuation. We found shipping the crystals transatlantic in the Blue Box resulted in no impact on data quality; we were able to solve comparable high-resolution structures from crystals regardless of whether they were shipped internationally or not. Interestingly, for KAT2B, we were able to solve structures of room-temperature datasets with improved resolution and data statistics compared to previously deposited cryocrystallography structures, albeit in a different crystal form. These results are promising in that they illustrate a way of transporting room-temperature crystal samples long-distance using standard, non-specialist commercial shipping. These experiments provide proof-of-concept that not only simplifies the setup of room-temperature X-ray diffraction experiments, but also demonstrates a viable workflow for the shipment of non-cryogenic protein crystals, which will allow more researchers from around the world to collect room-temperature crystallographic data at cutting-edge beamlines. To our knowledge, this is the first comprehensive study on the effects of shipping at ambient temperature on protein crystals, particularly with the goal of *in situ* data collection, although other groups have shipped crystals at ambient temperature.

## 2. Methods

### 2.1. Crystallization

Proteins were obtained as lyophilized powder and prepared for crystallization experiments as follows: Lysozyme from *Gallus gallus* (Hampton Research) was resuspended at 35 mg/mL in 20 mM sodium acetate, pH 4.6. Thaumatin from *Thaumatococcus danielii* (Sigma Aldrich) was resuspended at 50 mg/mL in distilled H_2_O. Thermolysin from *Geobacillus stearothermophilus* (Sigma Aldrich) was resuspended at 40 mg/mL in 45% dimethylsulfoxide (DMSO), 20 mM Tris HCl pH 8.0. Initial crystallization conditions for thaumatin and thermolysin were determined by a high-throughput (HT) screen performed at the National Crystallization Center at UB-HWI (Budziszewski *et al*., 2023, Lynch *et al*., 2023). The HT conditions were determined using the micro-batch under oil method, and yielded novel crystallization conditions for thermolysin. Preliminary hits were optimized in 96-well ARI Intelli-plate vapor diffusion plates using a Formulatrix Formulator to generate cocktails and a Formulatrix NT8 to set up drops. Further optimization was performed for the MiTeGen In Situ-1™ plates using the SPT Labtech Mosquito to dispense drops. Optimized crystallization conditions were as follows: Lysozyme was crystallized in 0.78 M sodium chloride and 0.1 M sodium acetate pH 4. Thaumatin was crystallized in 0.59 M di-ammonium tartrate. Thermolysin was crystallized in 0.20 M sodium di-hydrogen phosphate and 16.7% polyethylene glycol (PEG) 3350. Minimal optimization was required to translate crystallization conditions from the 96-well ARI Intelli-plate vapor diffusion plates to the MiTeGen In Situ-1™ plates.

For experimental setup at UB-HWI, optimized crystallization cocktails were dispensed into Greiner 96-well flat-bottom plates using the Formulatrix Formulator, with 35 microliters subsequently dispensed into MiTeGen In Situ-1™ plates using a multi-channel pipette. Two 200 nL drops were set in each 96-well In Situ-1™ plate using the SPT Labtech Mosquito at a 1:1 ratio of protein to cocktail. A single crystallization condition was used across all 96 wells for each protein sample. Plates were set up in duplicate for shipping. Additionally, Greiner 96-well flat-bottom plates with dispensed cocktails were shipped to Diamond Light Source along with lyophilized protein and buffer solution.

For samples set up at the Crystallisation Facility at Harwell Research Complex, lyophilized protein was prepared using the shipped buffers as stated above. Using a multi-channel pipette, 35 microliters of the optimized crystallization cocktails were dispensed from the shipped Greiner 96-well flat-bottom plate into MiTeGen In Situ-1™ plates. Two 200 nL drops (1:1 protein to cocktail) were dispensed into each 96-well In Situ-1™ plate using the SPT Labtech Mosquito. Care was taken to minimize procedural differences between plates set up at the two separate locations.

Purified KAT2B was thawed and centrifuged at 15,000 × rpm for 10 min. Initial hits were found in 0.1 M Tris pH 7.2, 8% PEG 8 K in Swiss Sci plates. These crystals were used to prepare a seed stock by crushing the proteins with a pipette tip, suspending in reservoir solution and vortexing for 60 s in the reservoir solution with ∼10 glass beads (1.0 mm diameter, BioSpec products). The condition was optimised for MiTeGen In Situ-1™ plates with 15 mg/ml and the final crystallization condition was 10% PEG 3350, 4.5% PEG 8000 and 0.1 M TRIS (HCl) pH 7.2 with a seed stock dilution of 1/10. The seeds were prepared from crystals grown in the final crystallization condition. The drop ratios were 0.2 µl protein, 0.07 µl reservoir solution, 0.03 µl seed stock. Crystals were grown using the sitting drop vapor diffusion method at 293 K using a SPT Labtech Mosquito robot and appeared within 24 h.

### 2.2. Shipping

The SSRL Blue Box is optimized for shipping crystals or other temperature-sensitive samples in microplates that conform to the SLAS standard footprint (https://www.slas.org/education/ansi-slas-microplate-standards/). The SSRL beamline 12-1 user community has reliably used the Blue Box since 2020 to ship SSRL *in situ* crystallization plates, compatible with an upgraded version of the Stanford Automated Mounter (Russi *et al*., 2016), for remote experiments with crystals maintained at controlled (non-cryogenic) temperature or controlled-humidity conditions. This includes successful commissioning shipments from Buffalo, NY to SSRL, Menlo Park, CA, and subsequent diffraction data collection.

The SSRL Blue Box consists of layers of impact isolating and thermally insulating materials (Figure 2). It features an outer layer of corrugated blue plastic lined with flexible foam that surrounds a rigid box made from polyurethane insulation. Adjacent to each inside wall of the polyurethane box is a removable plastic liner that contains a non-toxic, phase-changing gel. Finally, the core of the Blue Box contains three stacks of foam holders with cut-outs that secure up to six standard microplates, each with an additional small gel pack on top, or six of the taller SSRL crystallization plates.

**Figure 2.**
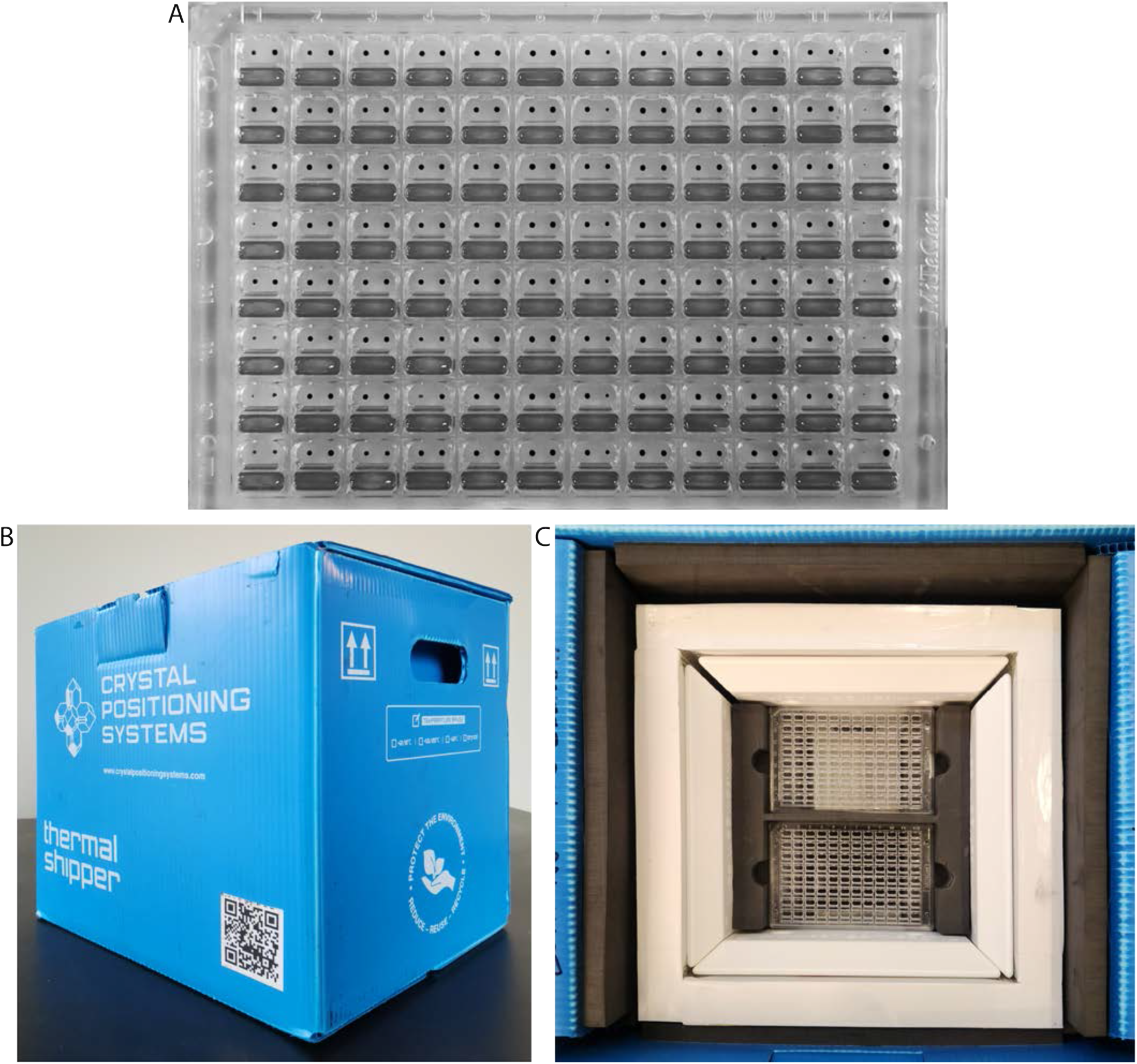
*In situ* crystal transport set up. A) Representative MiTeGen In Situ-1™ plate dispensed with food coloring in the reservoir and dye for the crystallization drop (shown in grayscale for visualization). B) SSRL Blue Box external and C) internal view with foam and insulation layers.

Before shipping, the box – either the entire box with the lid open or just the removable phase-changing gel liners – should be conditioned at 294 K (±2 K) for at least 48 hours. Once packed, crystallization plates inside the Blue Box can maintain temperatures between 288 and 298 K for up to several (4-7) days, depending on ambient weather conditions. In a similar manner, a second temperature option uses a different formulation of phase-changing gel to maintain temperatures between ±2 and ±8 degrees. The optimized Blue Box assembly has been commercialized by the company Crystal Positioning Systems and is sold directly or by distributors.

For our experiments, temperature packs were incubated at 296 K prior to packing samples. To address potential difficulties from sample handling or shipment delays while en route, duplicate plates were packed into two separate SSRL Blue Box Thermal Shippers and sent over two consecutive days via standard shipping option using a commercial shipping company. Samples cleared customs and arrived at DLS in 2-3 days from the shipment date.

In the return shipment of KAT2B, an Elitech LogEt 1 (www.elitechus.com) temperature data logger was included in the SSRL Blue Box, and set to record the temperature every 20 minutes upon activation.

### 2.3. Data collection, processing, and structure determination

Data were collected using the VMXi beamline at DLS. Crystals were selected and experimental parameters were applied using the SynchWeb interface to the ISPyB system (Delageniere *et al*., 2011) employed by DLS (Mikolajek *et al*., 2023). Individual crystals were selected within the interface and queued for data collection. Collection parameters were 10 µm x 10 µm beam size, X-ray energy 16 keV, an attenuated flux of approximately 1×10^12^ ph/sec, 0.0018s exposure, and a 60° rotation at 0.10° intervals for a total of 600 images with approximate X-ray diffraction weighted doses calculated using RADDOSE-3D (Zeldin *et al*., 2013) of ∼0.75 MGy. Datasets from multiple crystals auto-processed through the xia2.multiplex pipeline were selected for each experimental condition (Gildea *et al*., 2022). Each of these multicrystal datasets is derived from a number of individual datasets each measured from a single crystal. Several of these multicrystal datasets were then compared and their statistics averaged for comparison between crystallization set up location (US or UK). To achieve this, scaled, unmerged mtz files from the xia2.multiplex analysis for representative datasets were downloaded from ISPyB and processed using the following programs within the CCP4i2 software suite (https://www.ccp4.ac.uk).

For KAT2B, intensity-based clustering methods within xia2.multiplex were applied to the data as an additional test whether shipping the samples affected the properties of the crystal or protein structure. Intensity-based clustering methods within the DIALS software package are a way of identifying both systematic and random differences between datasets based on pair-wise correlation coefficients (Thompson *et al*., 2025). The shipped and non-shipped samples were analysed together using dials.correlation_matrix.

Lysozyme, thaumatin, and thermolysin structures were solved using MolRep (Vagin & Teplyakov, 2010) while KAT2B structures were solved using PHASER with a structure of KAT2B determined at 100 K as the search model (PDB 4NSQ (Shi *et al*., 2014, McCoy, 2007, McCoy *et al*., 2007)). Models were built and refined using Coot (Emsley & Cowtan, 2004) and Refmac5 (Murshudov *et al*., 2011). Additionally, statistics for 10 representative datasets from each condition were averaged for comparison (Tables S2, S7). The average age of crystals (lysozyme, thaumatin, thermolysin) at the time of data collection was approximately 16 days for those grown at UB-HWI, and 9 days for those grown at DLS. For KAT2B, shipped crystals were 27 days old at the time of data collection, while those that remained on-site at DLS were 7 days old.

## 3. Results

### 3.1. Experiments on Lysozyme, Thaumatin, and Thermolysin

#### 3.1.1. Setup and transportation of crystals

For the experiments using lysozyme, thaumatin, and thermolysin, entire 96-well plates were set up with a single protein and a single crystallization cocktail condition in order to generate a large number of similar crystals for each sample. Plates were imaged on the Formulatrix RockImager 1000 post-setup at UB-HWI for several days prior to shipment (Figure 3A-C). In an attempt to avoid shipping delays and mitigate potential rough handling of packages, duplicate crystallization plates were packaged in two separate SSRL Blue Boxes and shipped over two consecutive days to DLS in the United Kingdom. Plates arrived within 2-3 days of shipping.

**Figure 3.**
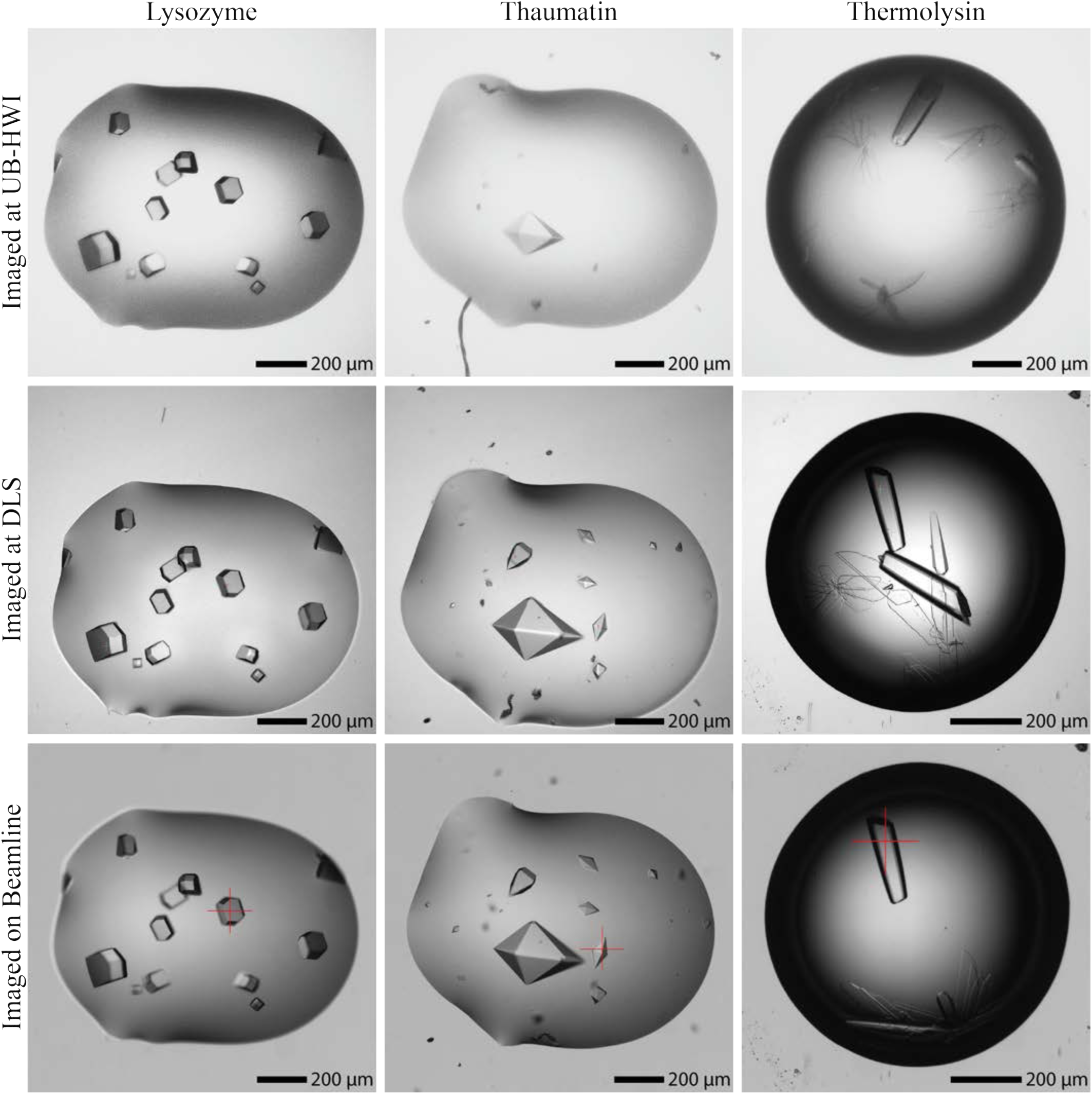
Images of crystals grown at UB-HWI in Buffalo, NY, imaged on UB-HWI’s Formulatrix Rock Imager 1000 (top row), imaged on DLS’s Formulatrix Rock Imager 1000 (middle row), and imaged on the VMXi beamline prior to data collection (bottom row).

Upon arrival at DLS, plates were imaged using the Rock Imager 1000 at VMXi (Figure 3D-F). Representative images of crystals on which data were collected are shown in Figure 3. Lysozyme and thaumatin crystals remained stationary in the plate during shipping, as seen in Figure 3. Visual inspection revealed no notable differences between crystal droplets from the two shipment boxes. Thermolysin crystals shifted during shipping (Figure 3F), and further shifted when the plates were rotated from horizontal to vertical for placement on the beamline (Figure 3I). Interestingly, the drops remained in place even when crystals within those drops moved. The crystal movement can be attributed to the 22.5% DMSO that is present in the crystallization cocktail and importantly is a feature of plate reorientation for data collection rather than being related to the crystal shipping process itself. Despite this movement, we were able to collect data on thermolysin crystals that either did not move or were re-marked for data collection in their new position following reimaging at the beamline. It can also be seen that thaumatin and thermolysin crystals continued to grow during the shipping process (Figure 3). Overall, no changes in crystal morphology or integrity were observed visually.

Crystallization reservoir solution, lyophilized protein and protein buffers were shipped to DLS to replicate the protein setup at UB-HWI. Identical plates were set up at the Crystallisation Facility at Harwell in order to make comparisons on data quality. Overall, crystals grown at DLS were of similar visual quality to those grown at UB-HWI.

#### 3.1.2. Room-temperature *in situ* X-ray diffraction experiments at VMXi

With the parameters used for data collection, minimal radiation damage was observed over the course of the experiment for all crystals (assessed by per image resolution plot). Each plate contained many crystals; therefore a few dozen datasets were collected for each experimental condition to ensure completeness from multicrystal datasets and to enable a robust comparison of data quality.

For data measured at VMXi, xia2.multiplex provides the best output data from multiple crystals when available, so we selected datasets processed through this pipeline for analysis (Table 1, Table S1). For lysozyme and thaumatin we compared 9 and 8 multi-crystal datasets respectively from UB-HWI and DLS grown crystals. Only 3 multi-crystal datasets were available for comparison for thermolysin due to the fact that crystals sank to the bottom of the well when the plate was rotated for placement on the beamline. While dozens of single crystal datasets were collected, auto processing only runs for multiple datasets collected from single crystallization drops. Subsequent to these experiments, VMXi now supports grouping multiple drops into a single sample group for auto processing. It is important to note that while crystallization and data collection conditions were replicated as closely as possible, crystals grown at UB-HWI and shipped to DLS were slightly older in age prior to data collection. Overall, the data quality of the crystals is effectively the same, regardless of whether they were shipped from UB-HWI to DLS or set up at DLS. The resolution, CC_1/2_ values, and R_pim_ are broadly consistent, with minimal variability that is likely due to slight differences from the crystals on each plate in crystal size and in the case of R_pim_ to differences in the number of contributing crystals to the datasets (R_pim_ is expected to decrease with greater multiplicity) (Table 1). The beamline geometry limits the circular resolution to 1.8 Å with data up to 1.5 Å available from the corners of the detector provided sufficient redundancy is present.

**Table 1.**
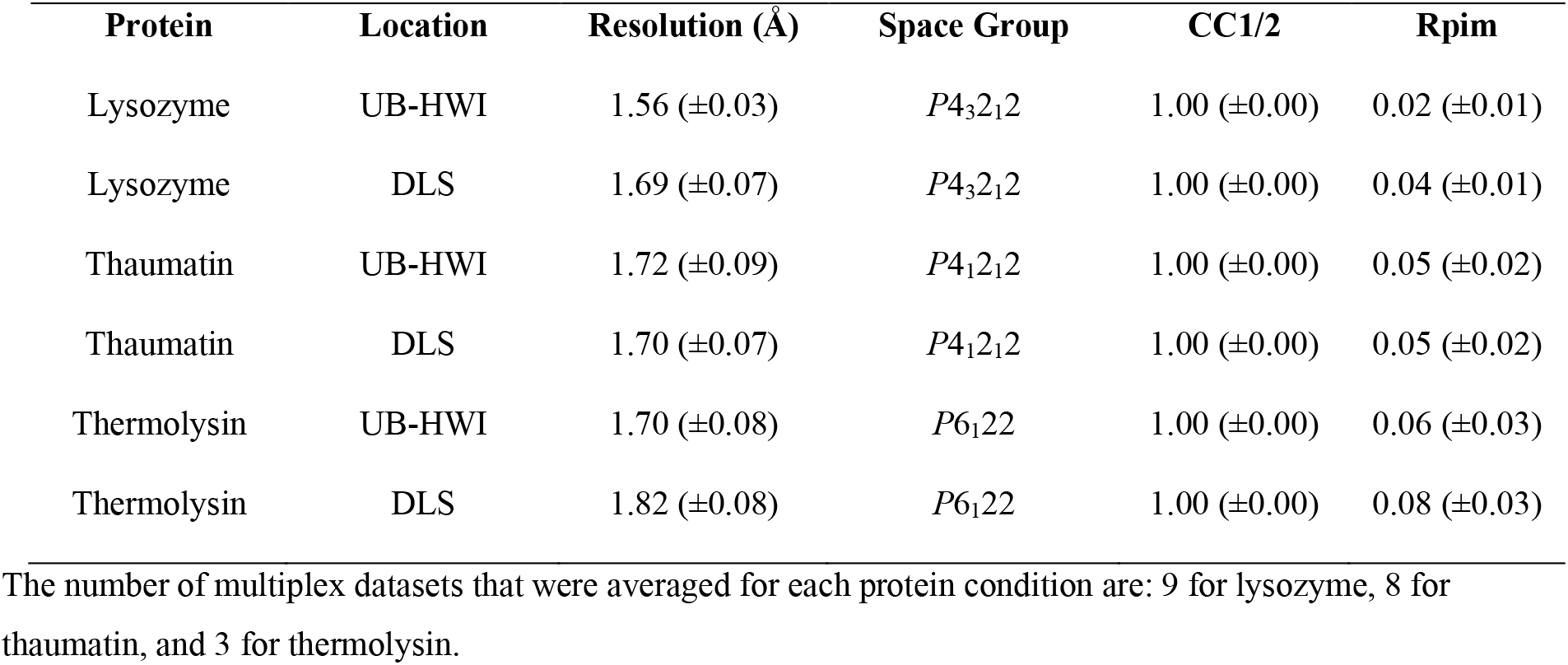
Comparison of average multiplex diffraction data quality.

Lastly, we compared the space groups and unit cells of the crystals to ensure there were no alterations due to transportation conditions, for example, dehydration of samples. Crystals grown at both locations indexed into their expected space groups; lysozyme in *P*4_3_2_1_2, thaumatin in *P*4_1_2_1_2, and thermolysin in *P*6_1_22, and the unit cell parameters are consistent with other non-cryogenic structures in the PDB. Additionally, in controlled dehydration experiments, unit cell contraction and relatively large changes in angles are observed (Sanchez-Weatherby *et al*., 2009, Russi *et al*., 2011), which we do not observe for crystals grown at either location suggesting that no significant dehydration has occurred during transport (Table S1).

We also performed an analysis of data statistics for 10 representative datasets from each experimental condition to determine the degree of variability across the experiment (Table S2). Overall, the results were similar to our multiplex analysis, with there being minimal variability in unit cell, resolution, CC_1/2_ values, and R_meas_, indicating consistency of crystal quality across conditions.

These data indicate transatlantic shipping of room-temperature protein crystals inside the Blue Box does not negatively affect the quality of diffraction data obtainable for the three crystal systems assessed. Crystals grown at UB-HWI and shipped long distance by air and road diffracted as well as those grown at DLS.

From these datasets, a representative dataset was chosen for each protein and setup location to process further and ultimately deposit coordinates and data into the PDB. These datasets generally had the best statistics from Table 1, in terms of resolution and R_pim_. Overall, the data processing statistics were very similar for crystals grown at each location for each of the 3 proteins (Tables S3-S4). Notably, all structures were solved to approximately 2 Å or lower resolution. We also analyzed the B-factors throughout the main chain and saw little variation between structures.

### 3.2. Experiments on KAT2B

To verify that our shipping protocols are viable for typical protein crystal samples, we carried out a similar shipping experiment using crystals of the human lysine acetyltransferase 2B (KAT2B). For this experiment, all crystallization plates were set up at DLS. Two plates were imaged at DLS and shipped to UB-HWI in the SSRL Blue Box. Upon arrival the plates were imaged, and then sent back to DLS in the SSRL Blue Box and imaged again (Figure 5). In the first shipment from DLS to UB-HWI, there is very minimal disruption of drop shape and crystal position, although the drop volume appears to be reduced slightly. During shipment back to DLS, there is a small change in drop shape (Figure 5). At the point of data collection upon DLS return, samples have been shipped internationally twice, and are 27 days old. To verify temperature variability in the SSRL Blue Box, we also included a temperature logger on the trip from UB-HWI to DLS which showed a reduction in temperature from 293.5 K to 289.2 K over the 64.8 hours of shipment, which is within the tested range of the SSRL Blue Box (Figure S1).

**Figure 4.**
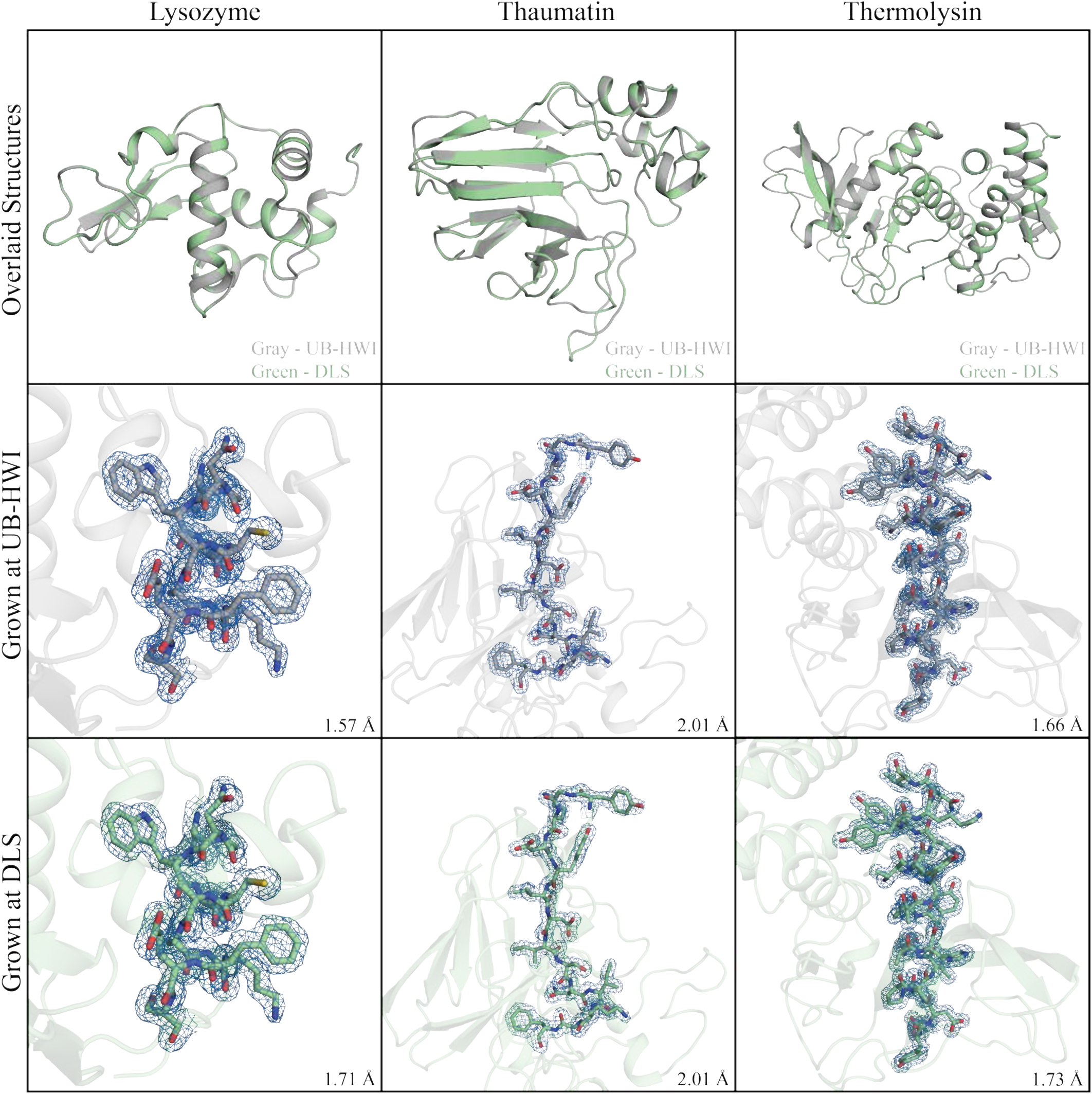
Representative structures with backbone alignment (top row), and representative view of side-chain electron density from 2Fo-Fc maps at 1.0 σ contour level for structures from crystals grown at UB-HWI (middle row) and at DLS (bottom row).

**Figure 5.**
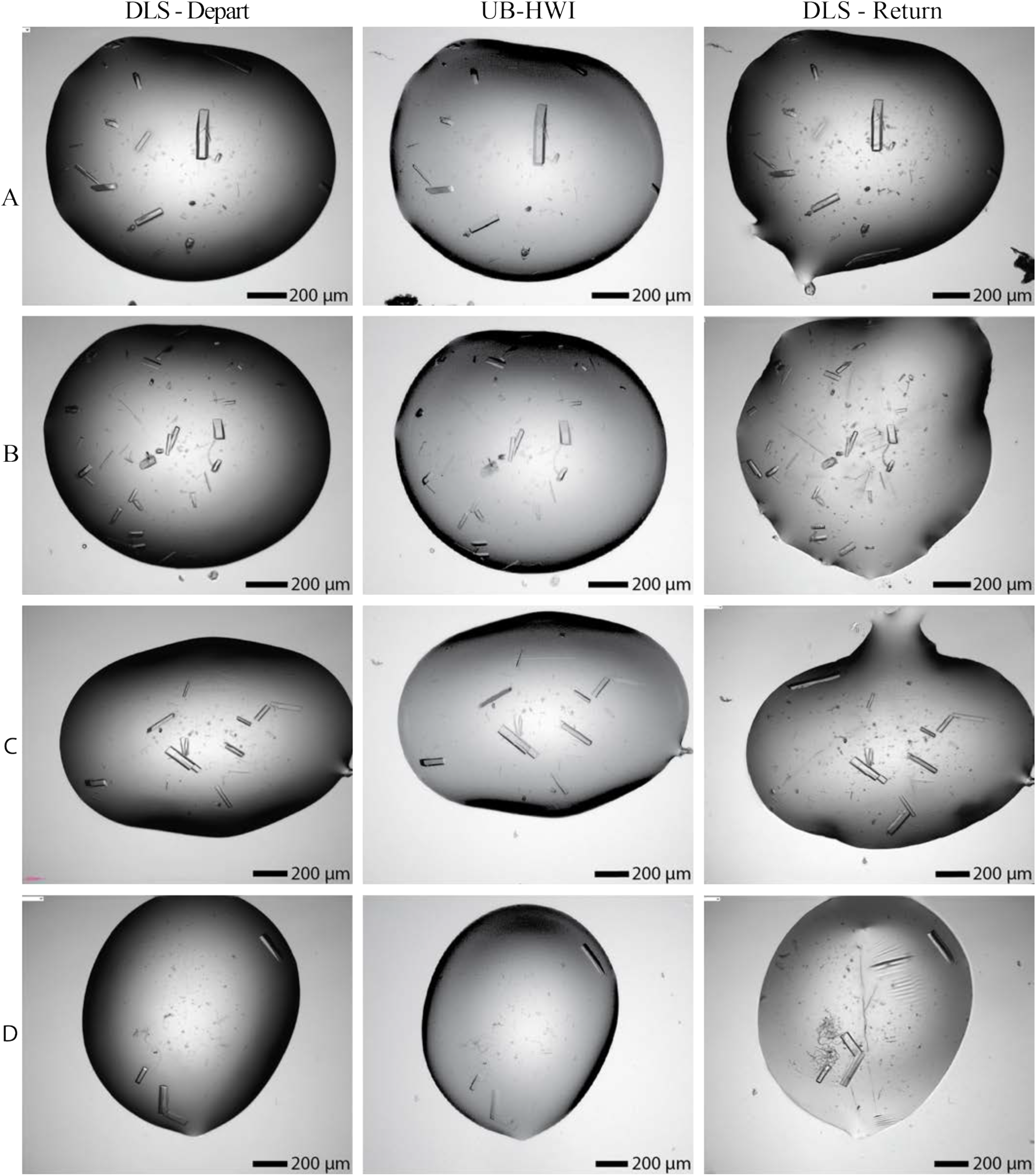
Images of four representative wells (A-D) of KAT2B crystals grown at DLS (left column), imaged at UB-HWI after shipment (middle column), and imaged after shipment back to DLS (right column).

Data were collected on both shipped and non-shipped KAT2B crystals that were each 27 days old having been set up in the same lab at the same time (Tables S6 and S7). Data from each group were pooled together, and 16 datasets for each used in our analysis. Structures from these multicrystal datasets were solved to 1.83 and 1.87 Å respectively (Figure 7). As with our other protein crystal samples, there was minimal variability in the merging and refinement statistics of final structures from shipped and non-shipped samples. Overall, these structures are comparable to the previously deposited structure in the Protein Data Bank PDB 4NSQ (Shi *et al*., 2014) determined at 100 K. Our VMXi structures are the first non-cryogenic structures of this protein to be determined. Notably, the RT crystals were indexed into a different space group to the previously published structure (PDB 4NSQ), *P*2_1_, and show an increase in resolution of ∼0.5 Å, likely due to the differing crystal forms, and there is electron density to model a previously unmodeled loop to varying extents between protein copies in the asymmetric unit for both shipped and non-shipped crystals.

**Figure 6.**
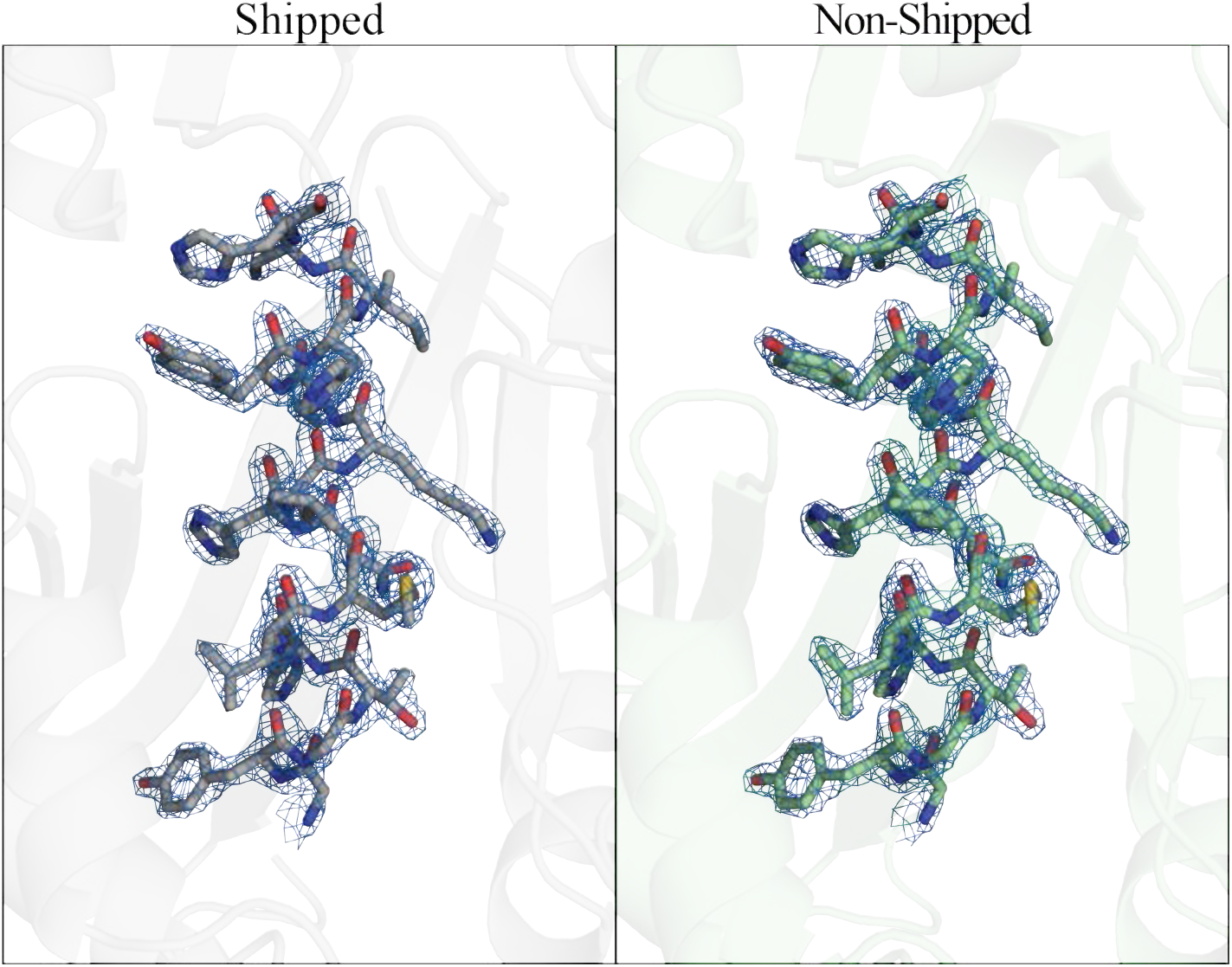
Representative view of side-chain electron density from *2Fo-Fc* maps at 1.0 σ contour level for structures from shipped (1.83 Å resolution) and non-shipped (1.87 Å resolution) crystals of KAT2B.

**Figure 7:**
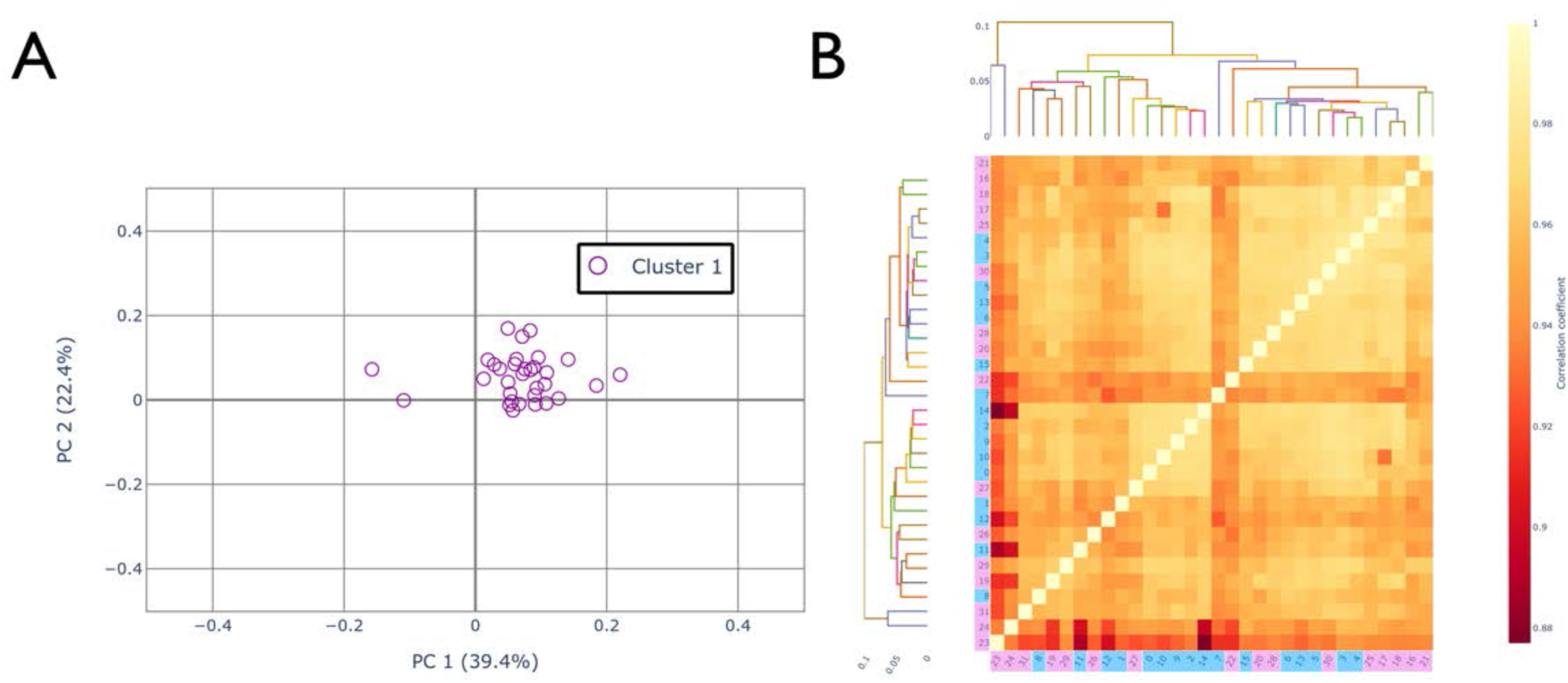
Intensity-based clustering analysis of shipped and non-shipped datasets for KAT2B. A) Coordinate plot of all datasets from the analysis in xia2.multiplex. All data appear to be in a single group indicating no systematic differences between shipped and non-shipped datasets. The X and Y axes correspond to the 2 primary principal components of the analysis, with the corresponding percentage of variance provided. B) Hierarchical cluster analysis of pair-wise correlation coefficients between all shipped and non-shipped datasets. The heatmap shows the pairwise CC values which all lie between 0.9 and 1. Pink datasets correspond to shipped crystals, and blue datasets correspond to not-shipped crystals. Given the shipped and non-shipped datasets do not group together, the dendrogram indicates no systematic difference in diffraction intensity or quality between these two experiments.

To verify no significant differences between the shipped and non-shipped samples, all data were further pooled together and analysed with DIALS (Gildea *et al*., 2022, Thompson *et al*., 2025). In the resulting plot (Figure 7A), each dataset is represented in coordinate space based on its similarity to other datasets; therefore, any systematic differences between datasets would result in well separated clusters of points. The OPTICS algorithm is used by DIALS to automatically identify high density regions to classify distinct subsets of data. In this case, all datasets were classified as belonging to a single cluster, meaning there were no statistically significant differences between shipped and non-shipped datasets. This is verified visually, as there is no systematic separation into distinct clusters. Using a more traditional approach, a hierarchical analysis of pair-wise correlation coefficients is also shown (correlation clustering in xia2.multiplex). It is clear from Figure 7B that all pairs of datasets correlate well (CC > 0.9), and there is no clear separation of shipped (pink) and non-shipped (blue) samples. Overall, these data further support use of our pipeline to facilitate routine, remote data collection of room temperature diffraction datasets where crystals are grown at the home lab and shipped for data collection.

## 4. Discussion

Data collection from crystals maintained at cryogenic temperatures has dominated X-ray crystallography for the past three decades, generating a wealth of structural data to populate the Protein Data Bank, contributing to the development of significant advancements in structural biology, and providing the training data for the remarkable success of AlphaFold (Jumper *et al*., 2021). It offers numerous advantages compared to room-temperature data collection, including increased tolerance to radiation damage, as well as enabling efficient transport of samples from home labs to beamline facilities.

However, cryogenic conditions can obscure biologically relevant conformations, in turn limiting insights into enzyme catalysis, ligand binding, and allostery. Recently, room-temperature crystallography is re-emerging as a desirable technique as it more closely represents physiological conditions. Major obstacles faced by room-temperature crystallography include the increased sensitivity to radiation damage, as well as the risk of crystal dehydration when crystals are removed from crystallization plates. These limitations highlight the need for improved experimental protocols.

The VMXi beamline at DLS specializes in room-temperature crystallography *in situ*, i.e., measurements of diffraction data directly from crystals within crystallization plates. This technique is advantageous over other methods of room-temperature data collection because the crystals are never removed from their crystallization environment, and therefore are not subjected to stress from manipulation such as fracturing or dehydration. Our experiment provided a challenging test case for shipping samples in plates; not only must crystals arrive in good condition, excessive movement of the crystallization drop within the plates would cause crystals to move out of the field of view of the imaging system required for *in situ* data collection. This represents a significantly more stringent test for shipping than simply shipping plates for crystal harvesting and mounting, where movement of a drop, provided it remains intact, is less of a problem. Our finding that drops remained in the field of view after transatlantic shipment demonstrates that the mail-in approach is reliable and enables a far wider community to access *in situ* data collection at ambient temperature.

We expected that the Blue Box would work well for transatlantic shipping because it has been used since 2020 to support a fully remote-access program for crystallography experiments at non-cryogenic temperatures and controlled humidity conditions at SSRL beam line 12-1, which is used routinely by fourteen user groups within the USA. While we are not the first to ship crystals at non-cryogenic temperatures, to our knowledge this study provides the first rigorous comparison of protein crystals grown and shipped at non-cryogenic conditions via transatlantic air freight by standard commercial courier, with those set up and imaged locally at a synchrotron source. While cryogenic shipping of protein crystals is well-established and widely used, the development of reliable, non-cryogenic shipping protocols is essential for expanding the accessibility of room-temperature crystallography to the global community of structural biologists.

Here, we show that high-resolution, room-temperature X-ray diffraction data can be collected from crystals shipped across the Atlantic Ocean. When comparing diffraction quality of crystals grown at UB-HWI and shipped to DLS to those grown at DLS, we observed no decrease in diffraction quality (Table 1, Table S6), and subsequently were able to determine high-quality structures of proteins from crystals grown at both facilities (Figure 4, Figure 6, Tables S1-6).

Data quality statistics for structural models from crystals grown at UB-HWI and shipped to DLS are comparable to those for the sample setup at DLS. It was hypothesized that thermal and mechanical stresses of transport by air using standard commercial shipment could result in a loss of diffraction quality, and changes to crystal integrity with regards to crystal packing and unit cell dimensions. Interestingly, virtually no changes in the unit cell of crystals grown at UB-HWI and transported to DLS compared to those grown at DLS were detected.

We have previously noted variability when reproducing crystals at different locations even with identical crystallization cocktail and protein preparation. This is likely due to factors outside of our control, such as environmental humidity, or other facility-specific factors. Nonetheless, these factors did not negatively impact diffraction data quality reported here.

The ability to measure X-ray diffraction data and determine structures from crystals with minimal additional handling is valuable for obtaining structural information from samples that are more difficult to study. Preparing crystals in the researcher’s laboratory and shipping for data collection decreases lab-to-lab variability (versus reproducing crystallization at the X-ray facility) and keeps sample preparation in the hands of the researcher all the way up to the step of measuring data. Importantly, these results illustrate an exciting new pipeline for growing and shipping crystals at room-temperature from the US to the VMXi beamline at DLS in the UK, and other cutting-edge beamlines around the world, increasing accessibility of room-temperature crystallography and empowering more studies on structural dynamics, fragment screening and ligand binding under noncryogenic conditions.

---

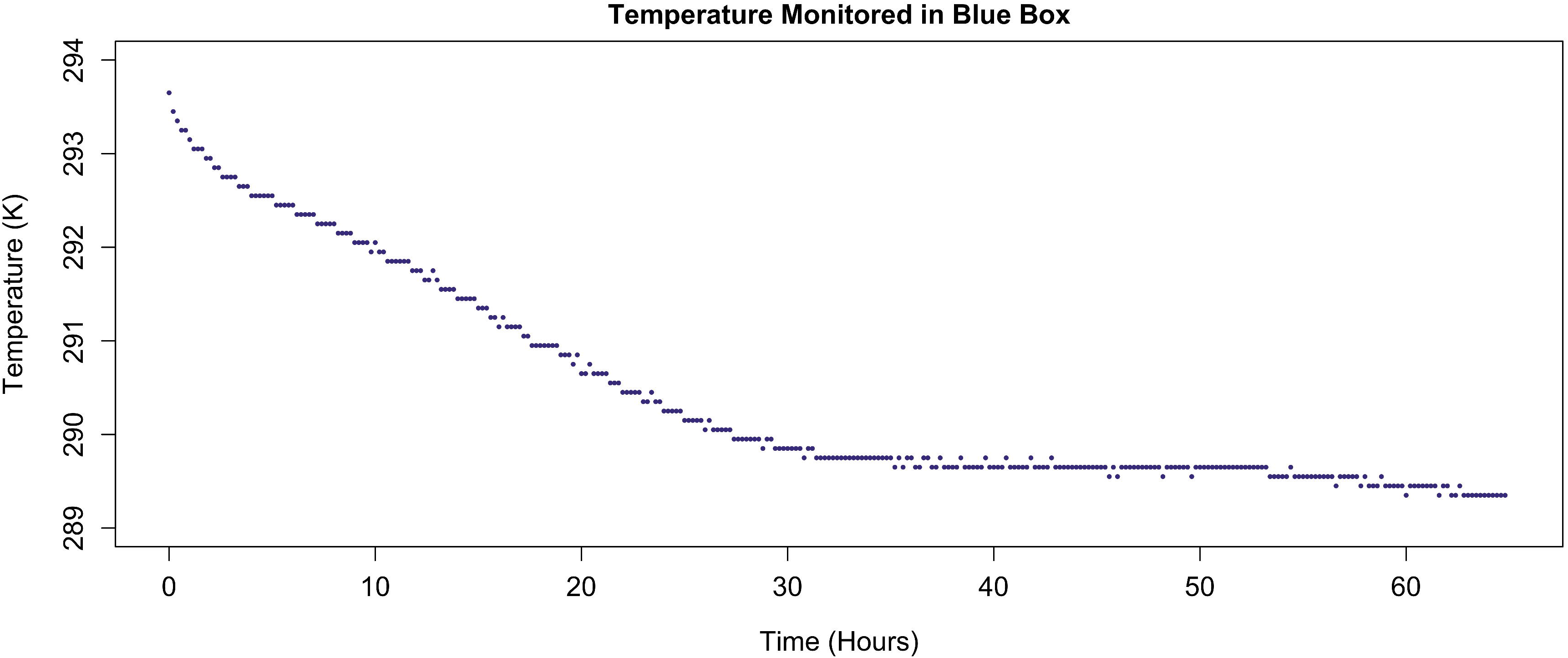

