## Supplementary Information for "An investigation on the feasibility of shipping protein crystals in plates for room temperature *in situ* data collection"

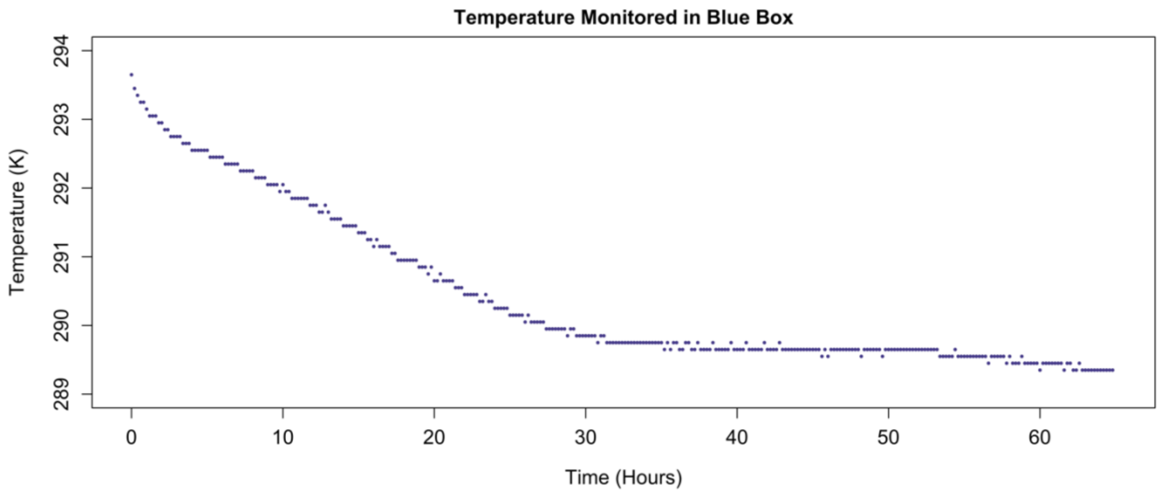


**Figure S1. Blue Box Temperature Log.** Using a Elitech LogEt1 placed in the SSRL BlueBox, the temperature of the KAT2B return shipment from UB-HWI to DLS was logged every 20 minutes over the course of the 64.8 hours of transit.

**Table S1** Averaged unit cell information for multiplex datasets

| Protein | Location | Unit cell (a,b,c (Å), α,β,γ(°)) |
|---|---|---|
| Lysozyme | UB-HWI | 78.65 (±0.01), 78.65 (±0.01), 37.69 (±0.01), 90, 90, 90 |
| Lysozyme | DLS | 78.67 (±0.01), 78.67 (±0.01), 37.70 (±0.06), 90, 90, 90 |
| Thaumatin | UB-HWI | 58.21 (±0.01), 58.21 (±0.01), 150.47 (±0.02), 90, 90, 90 |
| Thaumatin | DLS | 58.20 (±0.03), 58.20 (±0.03), 150.49 (±0.04), 90, 90, 90 |
| Thermolysin | UB-HWI | 93.06 (±0.04), 93.93 (±0.04), 130.53 (±0.07), 90, 90, 120 |
| Thermolysin | DLS | 93.09 (±0.02), 93.09 (±0.02), 130.59 (±0.02), 90, 90, 120 |

The number of multiplex multi-crystal datasets that were averaged for each protein condition in this analysis are: 9 for lysozyme, 8 for thaumatin, and 3 for thermolysin.

**Table S2** Analysis of representative individual datasets

| Protein | Setup Location | Resolution (Å) | Rmeas | CC1/2 | Completeness | Multiplicity |
|---|---|---|---|---|---|---|
| Lysozyme | UB-HWI | 1.63 (0.03) | 0.084 (0.016) | 0.997 (0.002) | 91.31 (2.81) | 3.74 (0.16) |
| Lysozyme | DLS | 1.76 (0.05) | 0.116 (0.020) | 0.997 (0.001) | 95.35 (2.54) | 4.08 (0.16) |
| Thaumatin | UB-HWI | 1.86 (0.08) | 0.196 (0.050) | 0.993 (0.004) | 97.72 (2.38) | 4.26 (0.16) |
| Thaumatin | DLS | 1.88 (0.08) | 0.210 (0.046) | 0.991 (0.003) | 98.94 (0.75) | 4.27 (0.16) |
| Thermolysin | UB-HWI | 1.77 (0.05) | 0.181 (0.030) | 0.994 (0.004) | 97.82 (1.50) | 6.11 (0.16) |
| Thermolysin | DLS | 1.80 (0.03) | 0.196 (0.024) | 0.995 (0.001) | 99.02 (0.87) | 6.17 (0.87) |

10 representative datasets were selected for each experimental condition, and their data averaged. Standard deviation is in parenthesis.

**Table S3** Structure solution and refinement data for lysozyme multiplex datasets

Values for the outer shell are given in parentheses.

| Protein (setup location) | Lysozyme (UB-HWI) | Lysozyme (DLS) |
|---|---|---|
| PDB ID | 9P12 | 9P13 |
| Number of crystals | 5 | 2 |
| Number of datasets | 5 | 2 |
| Resolution range (Å) | 55.63-1.57 (1.60-1.57) | 55.63-1.73 (1.74-1.71) |
| Unit cell a, b, c (Å) | 78.668, 78.668, 37.683 | 78.673, 78.673, 37.683 |
| α, β, γ (deg) | 90, 90, 90 | 90, 90, 90 |
| Space group | $P4_32_12$ | $P4_32_12$ |
| Completeness (%) | 99.8 (99.1) | 99.5 (95.5) |
| CC1/2 | 1.000 (1.000) | 0.998 (0.999) |
| I/σ | 1.17 (at 1.57 Å) | 1.24 (at 1.73 Å) |
| $R_{pim}$ | 0.02 | 0.04 |
| No. of reflections, working set | 17009 | 12788 |
| No. of reflections, test set | 839 | 616 |
| Final $R_{work}$ | 0.162 | 0.163 |
| Final $R_{free}$ | 0.203 | 0.207 |
| No. of non-H atoms | | |
| Protein | 1001 | 987 |
| Ligand | 1 | 1 |
| Water | 60 | 62 |
| R.m.s. deviations | | |
| Bonds (Å) | 0.015 | 0.015 |
| Angles (°) | 2.20 | 2.29 |
| Average $B$ factors (Å$^2$) | | |
| Protein | 23.89 | 25.05 |
| Ramachandran plot | | |
| Most favoured (%) | 99.2 | 99.2 |

| | | |
|---|---|---|
| Allowed (%) | 0 | 0 |

**Table S4** Structure solution and refinement data for thaumatin multiplex datasets

Values for the outer shell are given in parentheses.

| Protein (setup location) | Thaumatin (UB-HWI) | Thaumatin (DLS) |
|---|---|---|
| PDB ID | 9P15 | 9P14 |
| Number of crystals | 4 | 2 |
| Number of datasets | 4 | 2 |
| Resolution range (Å) | 54.27-2.01 (2.07-2.01) | 54.32-2.02 (2.07-2.01) |
| Unit cell a, b, c (Å) | 58.187, 58.187, 150.416 | 58.237, 58.237, 150.548 |
| α, β, γ (deg) | 90, 90, 90 | 90, 90, 90 |
| Space group | $P4_12_12$ | $P4_12_12$ |
| Completeness (%) | 99.8 (98.4) | 99.6 (99.2) |
| CC1/2 | 0.999 (1.000) | 0.997 (1.000) |
| I/σ | 1.14 (at 1.67 Å) | 1.15 (at 1.78 Å) |
| $R_{pim}$ | 0.05 | 0.06 |
| No. of reflections, working set | 17864 | 17583 |
| No. of reflections, test set | 844 | 832 |
| Final $R_{work}$ | 0.167 | 0.170 |
| Final $R_{free}$ | 0.204 | 0.210 |
| No. of non-H atoms | | |
| Protein | 1534 | 1556 |
| Ligand | 10 | 10 |
| Water | 206 | 206 |
| R.m.s. deviations | | |
| Bonds (Å) | 0.016 | 0.015 |
| Angles (°) | 2.39 | 2.43 |
| Average $B$ factors (Å$^2$) | | |

| | | |
|---|---|---|
| Protein | 23.3 | 22.9 |
| Ramachandran plot | | |
| Most favoured (%) | 98.53 | 98.04 |
| Allowed (%) | 1.47 | 1.96 |

**Table S5** Structure solution and refinement data for thermolysin multiplex datasets

Values for the outer shell are given in parentheses.

| Protein (setup location) | Thermolysin (UB-HWI) | Thermolysin (DLS) |
|---|---|---|
| PDB ID | 9P17 | 9P16 |
| Number of crystals | 2 | 2 |
| Number of datasets | 2 | 2 |
| Resolution range (Å) | 80.57-1.66 (1.69-1.66) | 80.60-1.73 (1.76-1.73) |
| Unit cell a, b, c (Å) | 93.000, 93.000, 130.400 | 93.073, 93.073, 130.561 |
| α, β, γ (deg) | 90, 90, 120 | 90, 90, 120 |
| Space group | $P6_122$ | $P6_122$ |
| Completeness (%) | 98.8 (91.0) | 100 (100) |
| CC1/2 | 0.998 (0.999) | 0.998 (1.000) |
| I/σ | 1.06 (at 1.66 Å) | 1.16 (at 1.73 Å) |
| $R_{pim}$ | 0.06 | 0.08 |
| No. of reflections, working set | 39541 | 35485 |
| No. of reflections, test set | 1991 | 1844 |
| Final $R_{work}$ | 0.147 | 0.147 |
| Final $R_{free}$ | 0.169 | 0.173 |
| No. of non-H atoms | | |
| Protein | 2673 | 2661 |
| Ligand | 11 | 10 |
| Water | 232 | 214 |
| R.m.s. deviations | | |

| | | |
|---|---|---|
| Bonds (Å) | 0.001 | 0.001 |
| Angles (°) | 1.778 | 1.757 |
| Average *B* factors ($Å^2$) | | |
| Protein | 19.72 | 19.79 |
| Ramachandran plot | | |
| Most favoured (%) | 93.35 | 93.67 |
| Allowed (%) | 4.43 | 4.43 |

**Table S6** KAT2B Multicrystal datasets

| Condition | Space Group | Resolution (Å) | Rpim | CC1/2 | Completeness | Multiplicity |
|---|---|---|---|---|---|---|
| Shipped | $P2_1$ | 60.18–1.83 (1.87–1.83) | 0.076 (2.123) | 0.998 (0.260) | 100.0 (100.0) | 15.6 (9.5) |
| Not Shipped | $P2_1$ | 87.71–1.87 (1.90–1.87) | 0.070 (1.159) | 0.984 (0.367) | 100.0 (100.0) | 16.8 (11.0) |

Values for the outer shell are given in parentheses.

**Table S7** Structure solution and refinement data for KAT2B multiplex datasets

Values for the outer shell are given in parentheses.

| Protein (setup location) | KAT2B (shipped) | KAT2B (non-shipped) |
|---|---|---|
| PDB ID | 35WL | 35WM |
| Number of crystals | 16 | 16 |
| Number of datasets | 16 | 16 |
| Resolution range (Å) | 60.18-1.83 (1.83-8.97) | 87.71-1.87 (1.87-8.97) |
| Unit cell a, b, c (Å) | 37.189, 87.738, 60.595 | 37.202, 87.711, 60.726 |
| α, β, γ (deg) | 90.000, 96.708, 90.000 | 90.000, 96.623, 90.000 |
| Space group | $P12_11$ | $P12_11$ |
| Completeness (%) | 100 (100) | 100 (100) |
| Multiplicity | 15.6 (9.5) | 16.8 (11.0) |
| CC1/2 | 0.998 (0.260) | 0.984 (0.367) |
| I/σ | 10.2 (0.7) | 10.3 (0.9) |
| $R_{pim}$ | 0.076 (2.123) | 0.070 (1.159) |
| No. of reflections, working set | 34117 | 32019 |
| No. of reflections, test set | 1726 | 1527 |
| Final $R_{work}$ | 0.158 | 0.162 |
| Final $R_{free}$ | 0.204 | 0.203 |
| No. of non-H atoms | | |
| Protein | 2577 | 2561 |
| Ligand | 0 | 0 |
| Water | 166 | 136 |
| R.m.s. deviations | | |
| Bonds (Å) | 0.0075 | 0.0082 |
| Angles (°) | 1.635 | 1.688 |
| Average B factors (Å2) | | |
| Protein | 32.24 | 33.07 |
| Ramachandran plot | | |

| | | |
|---|---|---|
| Most favoured (%) | 98.74 | 97.8 |
| Allowed (%) | 0.63 | 1.26 |
